# Mental Health Status of Adolescents During the COVID-19 Pandemic: A Cross-sectional Survey among the Bangladeshi Graduate Students at Dhaka City

**DOI:** 10.1101/2020.11.12.379487

**Authors:** Taha Husain, Mohammad Main Uddin, Saber Ahmed Chowdhury, Nazmul Ahsan Kalimullah

## Abstract

**Objectives:** To identify the level of Mental Health Status of Adolescents During the COVID-19 Pandemic among the Bangladeshi Graduate Student at Dhaka

**Method:** A cross-sectional survey was conducted with 330 students from different public and Private Universities in Dhaka, Bangladesh between April 01, 2020 and July 31, 2020 amid the COVID-19 lockdown period in Bangladesh. A standard, self-administered online questionnaire consisting of questions on socio-demographic variables, mental health status, as well as stress management sent to the respondents through social networking platforms. Data were analyzed using descriptive statistics, t-test, one-way ANOVA and correlation tests.

**Results:** The mean score of mental health status was 2.08 based on four points scale. They felt problem in decision making (3.04), in doing the things well (2.92), in enjoying normal day to day life (2.88), in playing a useful part in life (2.85), in doing their task (2.75), living in perfectly well and in good health (2.70). The respondents also developed a suicidal tendency (2.55), felt nervous in strung-up (2.24), took longer time to do things (2.14), felt tightness and pressure in head (2.12), and found themselves pressurized by various stuff (2.05). This study also found a significant positive relationship between mental health status and age, living with parents, and parents’ attitude. Finally, this study revealed that the respondents managed their stress by chatting with their friends, parents and siblings, and by sleeping.

**Conclusion:** Mental health status of adolescents was found moderate in this study. This study suggests further large-scale study including different socio-economic settings in order to figure out the real scenario of adolescents’ mental health status of the country during the pandemic.

## Introduction

Mental health is crucial to the overall well-being of individuals, communities, and societies because mental, physical and social functioning are interdependent (Devin and Farbod, 2016). Mental Health has now become a public health policy concerns for almost all countries around the world due to the extreme prevalence of SARS-COV-2 (Chevance et al., 2020; Szcześniak et al., 2021). Mental health is something differ from mental illness which is a condition that affects a person's thinking, feeling or mood (Herrman, Saxena, Moodie, and Walker, 2005). Prior to COVID-19 pandemic mental disorders accounted for 13% of the global problem of disease (Thapar, Pine, and Leckman, 2017), yet, the world was woefully unprepared to deal with the mental health impact of this pandemic (Diseases, 2020). Prior to this pandemic about 15 million people suffer from mental disorder of various categories (Islam and Biswas, 2015) and due to the lock-down people are suffering more fear of disease, anxiety and mental stress in Bangladesh (Mamun & Griffiths, 2020).

Adolescence is a critical period for mental, social and emotional development (Schwarz, 2009). The adolescent brain undergoes significant adaptability in light of physical, sexual, and intellectual challenges (Giedd, Keshavan and Paus, 2008; UNICEF, 2011). Several research study found prevalence of mental health disorder among the people at the adolescent and young age (Schwarz, 2009; Kalaiyarasan and Solomon, 2014; Kieling et al. 2011; Lee et al. 2014). Current pandemic makes this group of people more vulnerable to develop mental health disorder (Duan et al., 2020). For the adolescents, due to the lower incidence of infection and mortality than adults, medical professionals were less focusing on the unique clinical features of COVID-19 and mental health status of this age group (Ma et al., 2020).

Moreover, to prevent the outbreak of COVID-19, Bangladesh have been closed the academic institutions from the March 18, 2020, therefore, about 3.7 million students are staying at home (W. Ahmed, 2020). Educational institution plays an emergent role, not just in supplying educational resources to the student but also in offering students an opportunity to communicate with teachers and receive psychological counseling (Yeasmin et al., 2020). Evidence has shown from previous studies that the adolescents who experienced disasters might suffer from greater stress and trauma because of the lack of proper reactions and coping techniques (Roussos et al., 2005). Thus the COVID-19 pandemic may worsen existing mental health problems and lead to more cases among adolescents because of the unique combination of the public health crisis, social isolation, and economic recession (Golberstein et al., 2020).

Mental illness, across the spectrum of disorders, is both a direct cause of mortality and morbidity (WHO, 2006). Despite this the mental health care system in Bangladesh faces multifaceted challenges such as lack of mental health services from the skilled workforce, inadequate financial resource allocation and social stigma (Islam, & Biswas. 2015). Furthermore, pandemic stressors such as terror of infection, dissatisfaction and boredom, lack of knowledge, lack of personal space at home, and family's financial loss may have even more troublesome and enduring impacts on mental health of the adolescents (Brooks et al., 2020). Yet, there is no literature available in Bangladesh on the impact of COVID-19 pandemic on adolescent’s mental health. Thus, it becomes important to determine how extended university closures, stringent social distancing steps and the pandemic itself have impacts on the mental health status of graduate students in Bangladesh. Therefore, the general objective of this study is to identify the level of Mental Health Status of Adolescents During the COVID-19 Pandemic among the Bangladeshi Graduate Student at Dhaka. And the specific objectives are to determine the mental health status of the adolescents, to know the factors associated with mental health status of the adolescents, to examine the relationship between affecting factors and mental health status of adolescents.

## Methods

### Study design

A descriptive cross-sectional study was performed to identify the level of mental health status of the adolescents in the public and private universities in Dhaka, Bangladesh.

### Study Participants

The sample size of this study was estimated by using ‘G power analyses’. Following the conventional standard, the estimated sample size of this study was calculated with the acceptable level of significance (α=0.05), power (1-β=0.80), and effect size of (γ=30). Thus, the estimated sample size of this study was 300 and ten percent of samples were additionally added to avoid non-response rate and any missing data.

### Survey Instruments

General Health Questionnaire-28 (GHQ-28) was modified to use in this study. It was originally developed by Iveta NAGYOVA (2000) and validated by Shahab Rezaeian, (2016). This questionnaire had three parts. Part-I consisted with demographic data questionnaire (DDQ). Part-II consisted with mental health status related questionnaire (MHSRQ). The Likert scoring procedure (1= not at all, 2= no less than usual, 3= more than usual, 4= much more than usual) is applied and the total scale score ranges from 23 to 110. Part-III was involved with stress management related questionnaire (SMRQ). In this part, the Yes/No scoring procedure (0= yes, 1= no) is applied and the total scale score ranges from 06 to 110.

### Data collection Methods

The data collection of this study was performed between April 01, 2020 and June 31, 2020 among the graduate student at the Dhaka City. A self-administered online questionnaire was distributed to the respondents through social networking site in order to facilitate the survey efficiently.

### Data Analysis

Data analysis included descriptive statistics as well as inferential statistics approaches. Descriptive statistics for categorical variables included frequencies and proportions; means and standard deviations were utilized for continuous variables. Differences between categorical variables were assessed for significance using the chi-square test. Bivariate analysis–t-test, one-way ANOVA and correlation co-efficient test were performed in this study. The significance level is set at a p-value <0.05 here. Data analysis is performed using IBM SPSS Statistics for Windows (Version 23.0), IBM SPSS Amos (Version 23.0), and Microsoft Excel (Version 2016).

### Ethics Statement

The formal ethical approval of this research was received from the ethics committee of Dr. Wazed Research and Training Institute, Begum Rokeya University, Rangpur (Approval no: dwrti/000a5) and this research adhered to the Helsinki Declaration as updated in Fortaleza. Before their inclusion in this research, permission was received from respondents and their privacy was assured. Before continuing to the questionnaire, all the participants were told about the particular purpose of this study. Participants were only able to complete the survey through online once and were able to end the survey at any moment they chose. Anonymity and secrecy of the information were guaranteed. The competent authority has received formal ethical clearance for this study.

### Data Analysis and Findings of the Study

#### Socio-Demographic Characteristics of the Respondents

Table 1 shows the demographic characteristics of the study participants. The mean age of respondents was 21.26 years from the age from 20 to 24. Most of the respondents were male (61.8%), Muslim (93.6%), living with parents (92.7%), 85.5% of the students had a family income over 15,000 takas, 98.2% of the respondents had both their father and mother together, and most of the respondents’ parents showed positive attitude to them.

**Table 1:** Socio-Demographic Characteristics of the respondents (n= 330)

| Variables |  | N (%) | M(SD) |
| --- | --- | --- | --- |
| Age |  |  | 21.26(.61) |
| Gender | Male | 204 (61.8) |  |
|  | Female | 126 (38.2) |  |
| Religion | Islam | 309 (93.6) |  |
|  | Hindu | 21 (6.4) |  |
| Nature of living arrangement | With parents | 306 (92.7) |  |
|  | Others | 24 (7.3) |  |
| Monthly Family Income (in Taka) | <15000 | 48 (14.5) |  |
|  | >15000 | 282 (85.5) |  |
| Parents marital status | Married | 324 (98.2) |  |
|  | Separated | 06 (1.8) |  |
| Parents' attitude to Students | Very positive | 264 (80) |  |
|  | Little bit positive | 48 (14.5) |  |
|  | Little bit negative | 18 (5.5) |  |

**Table 2:**
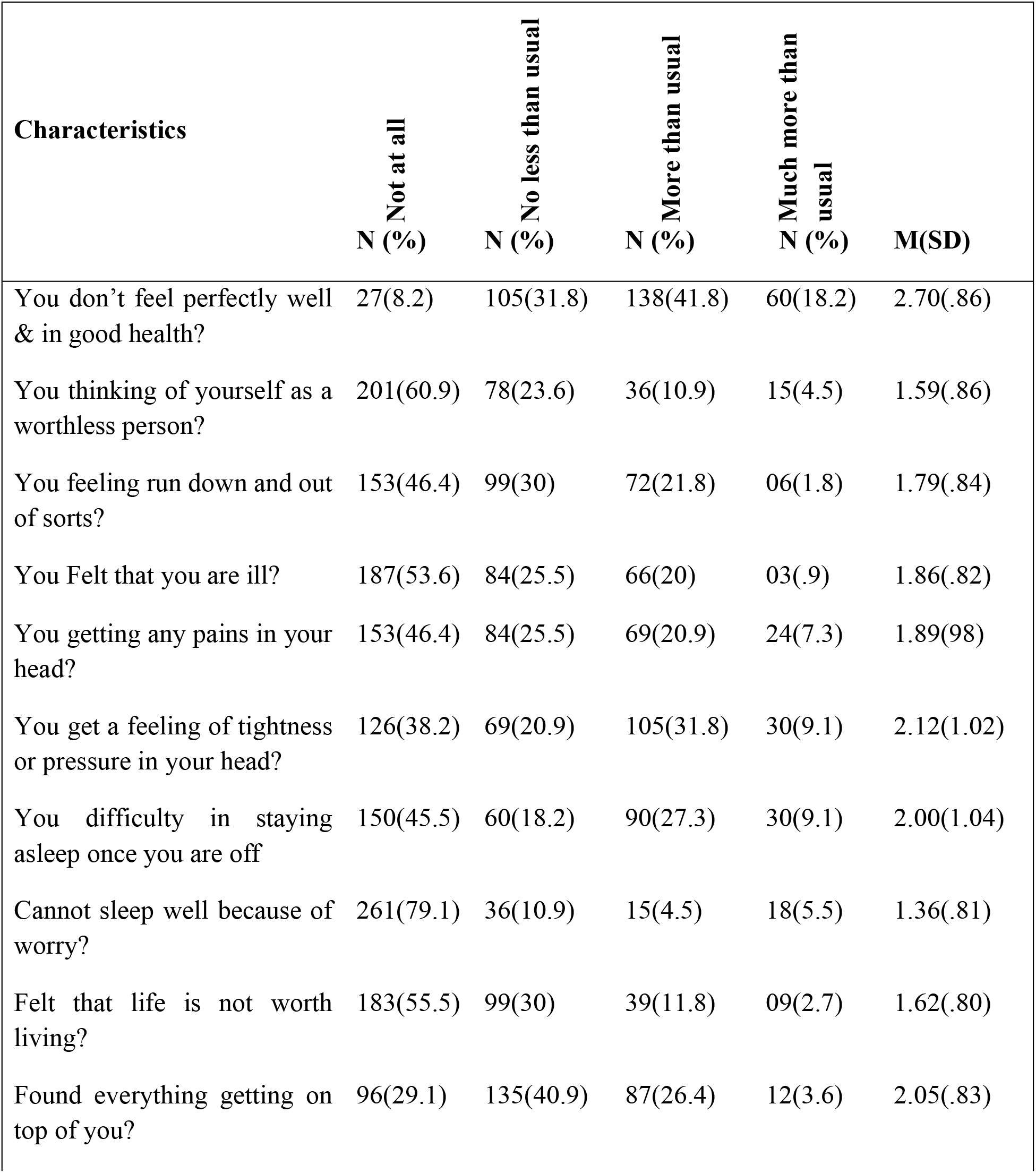
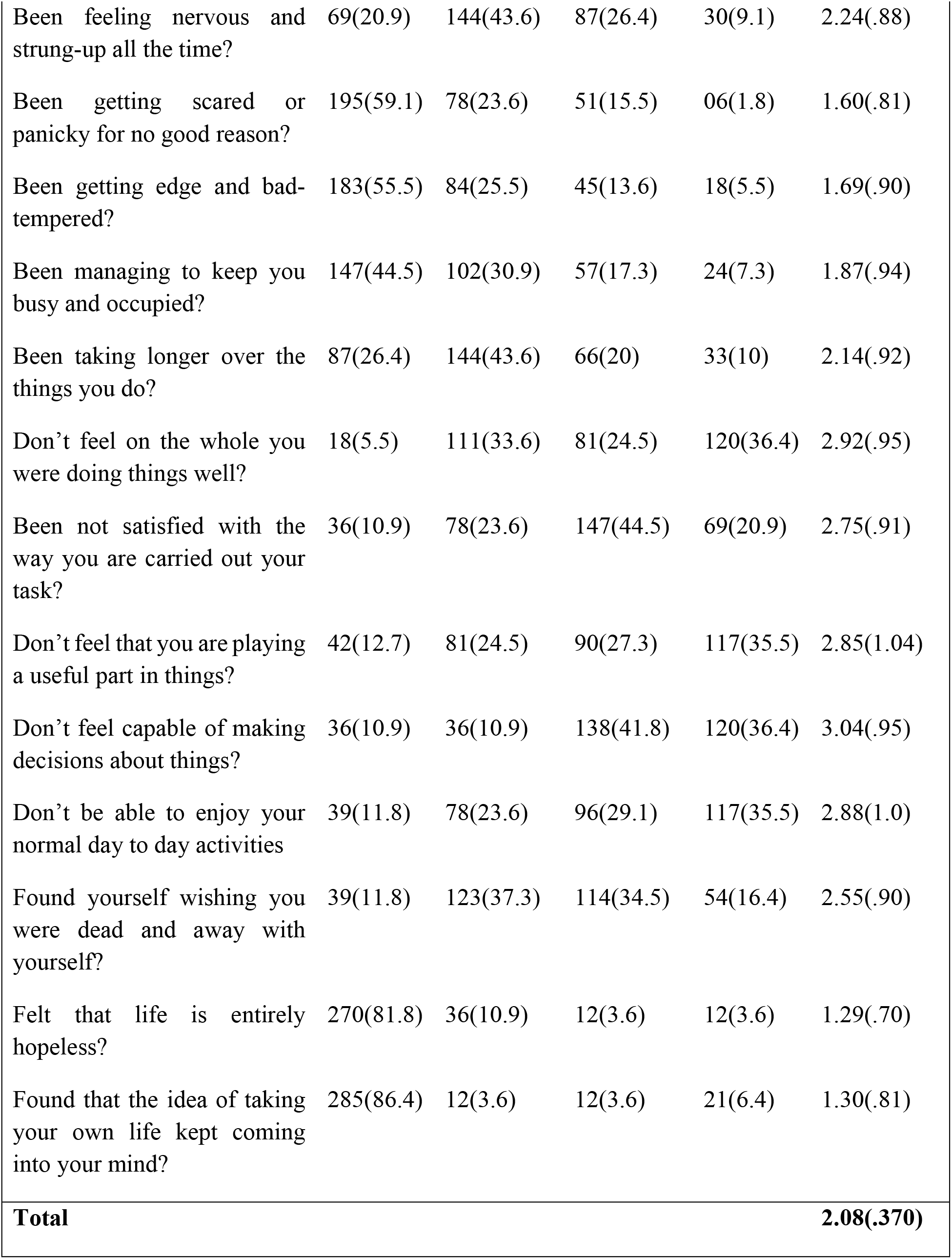
Mental health status of adolescents graduation level students during the lock-down period in Bangladesh (n=330)

#### Mental health status characteristics of adolescents during COVID-19

Table 3 shows the mental health status related characteristics of the study participants. By assessing mental health status related questionnaire 23 (MHSRQ-23) it depicts that the total scale score was 2.08 (.37%) which means for overall results the respondents had poorer psychological well-being. Among the 23 scales 11 scales was found significant. Students who didn’t feel capable of making decisions about their future amid the COVID-19 the total scale score were high 3.04 (.95%) than others. On the other side, participants who felt that they are not doing the things well (2.92), felt that they don’t able to enjoy normal day to day life (2.88), felt that they are not playing a useful part in life (2.85), felt they are not satisfied the way they are taking their task (2.75), don’t felt they are perfectly well and in good health (2.70), wished that they were dead and away with themselves (2.55), felt nervous and strung-up (2.24), took longer time to do things (2.14), felt tightness and pressure in head (2.12), and found everything get on top of them (2.05). on the contrary the respondents who with their life was entirely hopeless during the lockdown period of COVID-19 had a total scale score of 1.29 which was lower than others. Performers who answered that the idea of taking own life kept coming into their minds had the total scale score of 1.30 (.81) which was also lower than others still it is a concerning issue. Respondents whose worry kept them from sleeping had the total scale score of 1.36 (.81) lower than 3.04 (.95%). Thus it constitutes that sleep may reduce the mental anxiety and stress.

**Table 3:** Relationship between Demographic related characteristic and Mental Health Status Related Questionnaire (MHSRQ-23) (n=330)

| **Variables** |  | **M(SD)** | **t/f/r(p)** |
| --- | --- | --- | --- |
| **Age** |  | 17.26(.61) | -.194(.042) |
| **Gender** | Female | 2.9(.36) | .004(.997) |
|  | Male | 1.9(.37) |  |
| **Religion** | Islam | 2.9(.36) | .861(.391) |
|  | Hindu | 1.9(.49) |  |
| **Nature of living arrangement** | With parents | 2.6(.36) | -2.385(.019) |
|  | Others | 2.3(.39) |  |
| **Monthly Family Income in BDT** | <15000 | 2.0(.29) | -.085(.932) |
|  | >15000 | 2.0(.38) |  |
| **Parent's marital status** | Married | 2.0(.36) | -1.867(.065) |
|  | Other | 2.5(.00) |  |
| **Parents' Attitude** | Very Positive | 2.04(.37) | 3.354(.039) |
|  | Little bit positive | 2.27(.36) |  |
|  | Little bit negative | 2.26(.19) |  |
| **Close Friends** | No one | 2.3(.25) | 2.323(.103) |
|  | Less than three | 2.0(.38) |  |
|  | More than Three | 2.0(.36) |  |

#### Relationship between demographic characteristic and mental health status

Table 4 illustrates the relationship between demographic and the level of mental health status related questionnaire (MHSRQ-23). Accordingly, the findings revealed that there is a significant positive relationship between mental health status and age, living with parents, and parents’ attitude with the P-value of 0.042, 0.019, and 0.039 respectively. And categories such gender, religion, income, parents’ marital status, and having more friends’ has positive but insignificant relations with the P-value of (p> 0.05).

**Table 4:**
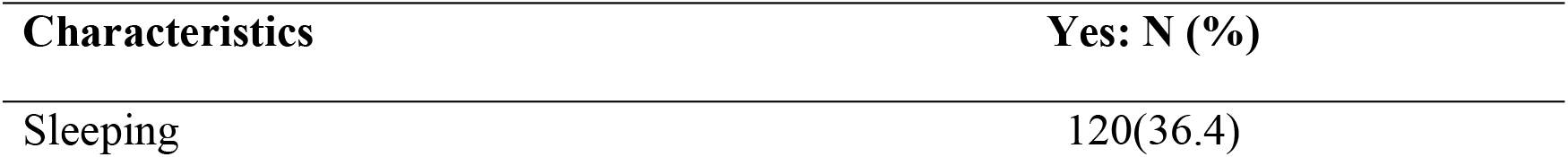
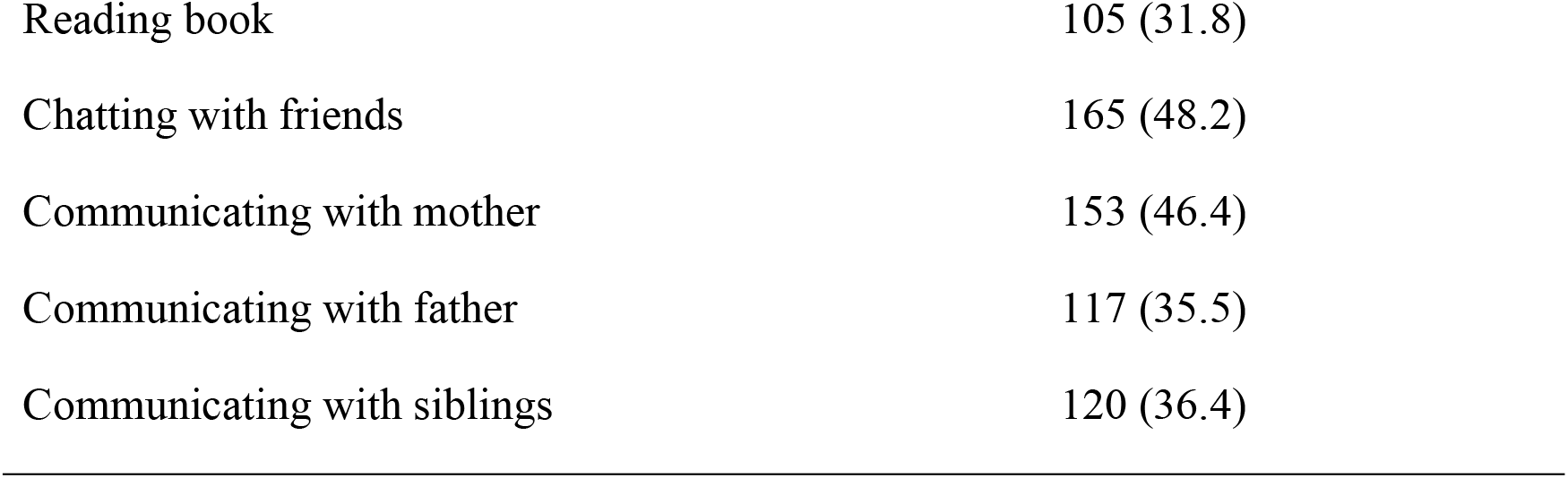
Stress management strategies of Adolescents (n=330)

#### Stress management strategies of adolescents during COVID-19

Table 2 shows the distribution of stress management characteristics of the study participants. Most of performers (48.2%) answered that they managed their stress by chatting with their friends, 46.4% by communicating with their mother, 36.4% by sleeping and communicating with siblings, 35.5% by communicating with their father, and 31.8% by reading books.

## Discussion

This study examined mental health status, relationship of mental health status with socio-demographic factors and the strategies of stress management of the adolescent respondents from graduation level students during the lock-down period in Bangladesh. This study found that adolescents people in Bangladesh developed significant mental health problem. They felt problem in decision making, doing the things in well manner, in enjoying normal day to day life, in playing a useful part in life, in doing their task, living in perfectly well and in good health. The respondents also developed a suicidal tendency, felt nervous in strung-up, took longer time to do things, felt tightness and pressure in head, and found themselves pressurized by various stuff.

Past studies in the context of china, Iran and many other countries of the world found the impact of COVID-19 on the respondents’ mental health (Gao et al., 2020; Pieh et al., 2020; Rajkumar, 2020; Zandifar & Badrfam, 2020). Literatures also found significant mental health problem for the adolescent and the students in the context of international Chinese students affected by the COVID-19 outbreak (Zhai & Du, 2020). It is also evidence from the Chinese college students that the COVID-19 isolation policies affected young people’s mental health (Chen et al., 2020). On the other side, based on the results of the univariate analysis, 0.95% of the students were incapable of making decision and were in a negatively associated with mental health status. Most of the students answered that they were incapable to make decision because in Bangladesh most of the students were worried about their future during the lock-down period (Bodrud-Doza et al., 2020). Due to this, students were not able to make their own decisions.

This study also found that respondents often develop anxiety and stress related mental health complexities due to the severe lock-down. Past studies in the context of Bangladesh, India found the same results along with other economic impact of the countrywide massive lock-down (Banerjee, 2020; Bhuiyan et al., 2020). Some other studies investigated impact of COVID-19 lockdown on agriculture, women domestic workers, marital life, general population and their habits, sleep-wake schedule and associated lifestyle related behavior in India (Ghosh et al., 2020; Kumar & Dwivedi, 2020; Maiti et al., 2020; Rawal et al., 2020; Sinha et al., 2020). These studies supported our findings that the common people around the world were imposed to stay home and did not perform their daily activities in a smother way. Thus, as shown earlier, that the consequences of COVID-19 including disruption in daily life, risk of infection, sense of confinement, inadequate supplies, fear, and anxiety associated with the virus can lead to detrimental effects at individuals and societal levels (Ahorsu et al., 2020; Arslan et al., 2020).

The findings of this study revealed that adolescent people in Bangladesh developed suicidal tendency during the COVID-19 pandemic. This finding echoes the study of (M. Z. Ahmed et al., 2020) in the context of China which has been revealed that a great anxiety, depression, alcohol use disorder of young aged people associated to lower mental wellbeing during the COVID-19 epidemic. Another study in Bangladesh found that being female, being divorced, and having no child were emerged as independent predictors for suicidality(Mamun et al., 2020). A study in Turkey also consistent with the findings of this study which depicts that COVID-19 perceived risk increases death distress and reduces happiness (Yıldırım & Güler, 2021).

This study found that people already felt pressure, nervous, and loss of productivity. It could be the impact of the implementations of these virus prevention and control measures have been effective to protect people against COVID-19, the risks of a second wave and a new peak of infections may cause uncertainties among people around the world (Yıldırım & Güler, 2021). Past studies have same consistent results in the context of Hong Kong which showed the worry and disruption of daily life due to the COVID-19 pandemic (Kwok et al., 2020). Other studies have also reported experience of various mental health difficulties such as depression, fear, anxiety, boredom, worry, sadness, sense of being trapped, feelings of insecurity, loneliness, and helplessness during the pandemic (Xiao, 2020).

This study also found a significant positive relationship between mental health status and age, living with parents, and parents’ attitude towards them. Consisting with the findings of this study other researches in India indicated that the adolescents from the high income group had good mental health status compared to students belonging to low income group families (Sharma, 2013). This study could not show any association between having divorced parents and a risk of self-rated mental health of the adolescent. The study of Wille et al. (2008) contrasted with this finding and revealed that the children who in high risk family get stressed.

Finally, this study revealed that the respondents managed their stress by chatting with their friends, parents and siblings, and by sleeping. Therefore, this study highlights the finding that adolescents manage of their stress by sleeping was 36.4% in a negative relationship with mental health status, and they manage their stress by reading books was 31.8%, Chatting with friends was 48.2% in a positive relationship with mental health status. So, the role of stress management in mental health and risk behaviors is particularly important in this age group.

The findings of this study cannot be generalized as the sample size was limited due to the current situation, it was not possible to collect samples on a large scale. The sample size could be larger in order to provide more powerful results. Another limitation was the exclusive use of self-report measures, a strategy often associated with method variance. The outcome of the study depended on the participants’ honesty and cooperation in answering the questions.

Considering health threats, a face-to-face interview was avoided whereas compared to face-to-face interviews, self-reporting has certain limitations.

## Conclusion

This study revealed that the mental health status of adolescents in Bangladesh during the lockdown period was moderate. This study finds some socio-demographic variables such as age and living status which are fostering mental health problem during the lockdown period in Bangladesh among the adolescent graduate participants. This study also finds the stress management strategies such as sleep and chatting to their near people. Also, the study was conducted only among the graduate students exclusively in an urban setting. Further studies can be carried out to study the relationship of all three attributes of Socio-economic status such as education, occupation and income on mental health among the all population with a larger sample and setting will be organized in urban and rural area as well. Also, the pattern of family relationship can be studied for adolescent boys and girls belonging to high socio-economic status and low socio-economic status separately.

